# Alternatively Folded State of IgG4: Acid-Induced Compaction is Common between IgG1 and IgG4

**DOI:** 10.1101/2025.02.08.637241

**Authors:** Hiroshi Imamura, Shinya Honda

## Abstract

Immunoglobulin G1 (IgG1) antibodies undergo denaturation in acidic conditions, resulting in an alternatively folded state (AFS). The AFS structure is more compact than the native state. However, the prevalence of AFS in other subclasses remains largely unexplored. This study provides evidence that humanized IgG4 can also adopt AFS structure, as demonstrated through size-exclusion chromatography coupled with small-angle X-ray scattering (SEC-SAXS) analysis. These findings suggest that the anomalous compaction of immunoglobulins G (IgGs) is resilient to variations in sequence and structure among subclasses.

## Introduction

Antibodies have become increasingly indispensable in biological, medical, and pharmaceutical sciences due to their molecular diversity and specific functionality. At the same time, their molecular characteristics, including (un)folding-related physical degradation, have been widely recognized as a concern, particularly for the safe production of antibodies in the biopharmaceutical industry[1,2]. One of the longstanding issues is the non-native but stable state of an antibody in addition to its native state. This state was first identified in 1991 and is referred to as an alternatively folded state (AFS)[3]. The AFS is typically induced by acidic conditions[3,4] but can also arise through heating[5]. As a thermodynamically stable state, the AFS molecule exhibits cooperative conformational transitions in the presence of a denaturant[3]. AFS species are prone to aggregation[3,4]; this poses challenges for industrial purification processes[6,7]. But, AFS antibodies may hold biological promise for potential oral administration due to their resistance to enzymatic digestion[8].

The elucidation of the AFS structures of antibodies has long been hindered, and even the relationship between AFS and oligomerization remains controversial. In 2023[9], we demonstrated that a monomeric antibody (humanized IgG1) can adopt AFS using size-exclusion chromatography (SEC) coupled with small-angle X-ray scattering (SAXS) (SEC-SAXS). A notable finding regarding IgG1 AFS was its well-folded structure (albeit denatured due to the loss of binding function[10]) with a reduction in its size (approximately 25% reduction) compared with that of its native form[9]. We postulated that the non-native domain-domain interactions are necessary for the anomalously compact structure, irrespective of whether these interactions are intramolecular or intermolecular. However, the molecular or sequence requirements for AFS formation remain unknown. It is unclear whether the capability to be AFS is unique to a specific antibody sequence or subclass. To date, all reports on antibody AFS have focused on IgG1[3,4], except for heat-induced oligomerization of the crystallizable fragment (Fc) of IgG4[5]. This study does not focus on the oligomeric state but instead examines the monomeric state. Our previous investigation of two IgG1 sequences suggested that the formation of monomeric AFS is insensitive to differences in the variable region sequence[9].

To further address this question, the present study targeted the IgG4 subclass and examined its AFS formation capability. The primary difference between humanized IgG1 and IgG4 lies in the arrangement of disulfide bonds and the length of the hinge region (Figure 1). Additionally, there is a minor difference in their constant region sequences. Although both subclasses share 12 domains and a Y-shaped structure, they differ in static and dynamic conformations[11]; this contributes to differences in FcγRI binding capacities[12]. Comparing IgG4 with IgG1 allows us to examine the universality of antibody AFS and provides insights into the molecular nature of the AFS structure. To this end, we applied the SEC-SAXS approach to an IgG4 monoclonal antibody.

**Figure 1.**
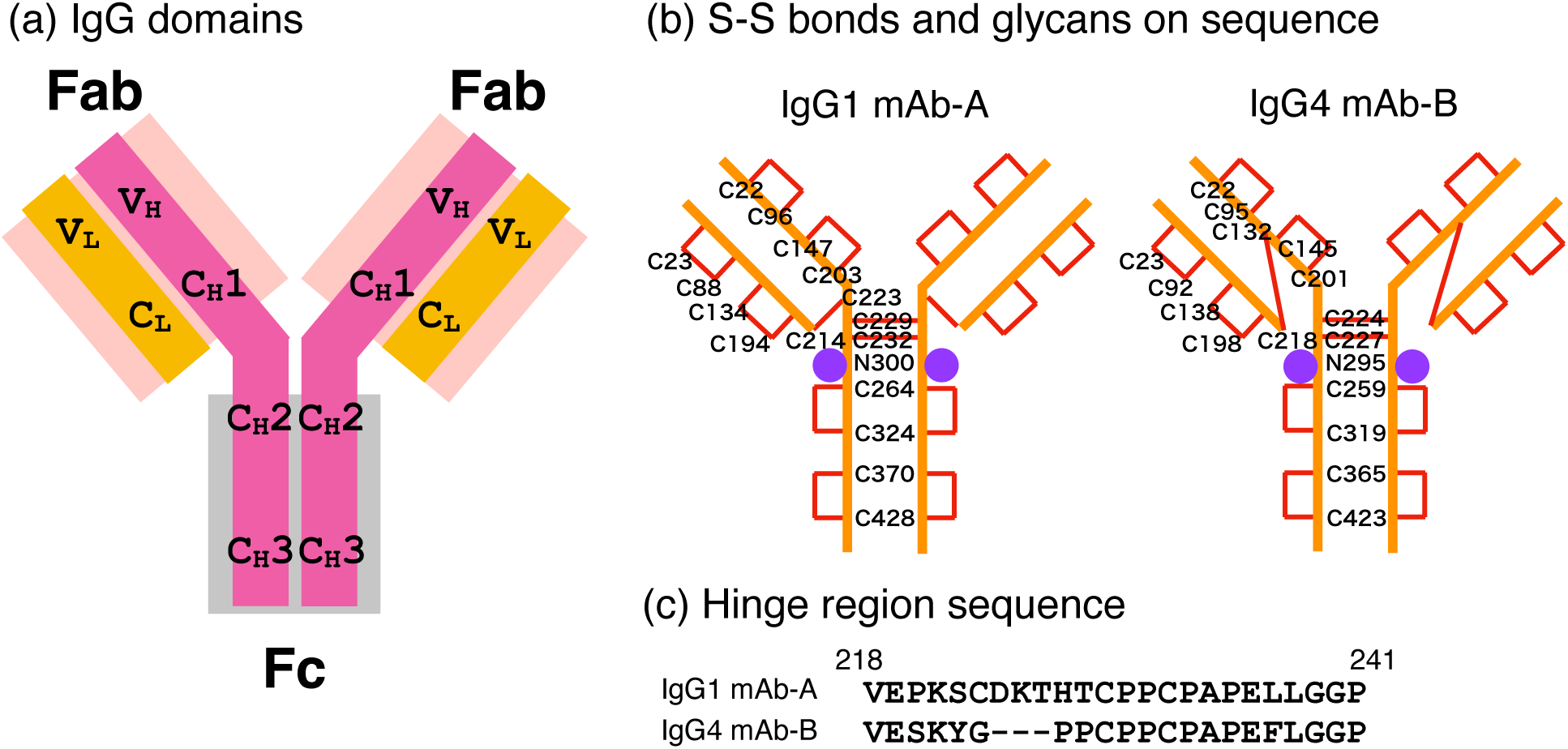
Structure of humanized IgG1 and IgG4. (a) IgG domains. (b) Disulfide (S-S) bonds (red line) and glycans (filled purple circle) in the sequences of IgG1 mAb-A and IgG4 mAb-B. (c) Hinge region sequence. Fc refers to the crystallizable fragment, and Fab refers to the antigen-binding fragment.

## Materials and Methods

We used a recombinant IgG4 monoclonal antibody, Lebrikizumab, which is commercially available as Ebglyss (Eli Lilly and Company, IN, USA) and referred to as mAb-B in this paper. According to the interview form[13], it is described as “composed of complementarity-determining regions derived from mouse anti-human interleukin-13 monoclonal antibody and framework regions and constant regions derived from human IgG4, whose amino acid residue at position 226 in the H-chain is substituted by Pro. Lebrikizumab is a glycoprotein (molecular weight: ca. 148,000) composed of 2 H-chains (*γ*4-chains) consisting of 445 amino acid residues each and 2 L-chains (*κ*-chains) consisting of 218 amino acid residues each.” For comparison, we used a humanized IgG1 monoclonal antibody (148 kDa), Trastuzumab, which is commercially available as Herceptin (Chugai Pharmaceutical Co., Ltd., Tokyo, Japan) and referred to as mAb-A in this paper.

### Size-exclusion chromatography (SEC)

Size-exclusion chromatography was performed using a Prominence HPLC system (Shimadzu Co., Kyoto, Japan) equipped with a Superdex 200^TM^ Increase 10/300 GL column (300 mm length × 10 mm internal diameter; Cytiva Sweden AB, Sweden, lot 10299331). Ultraviolet absorption of the eluate was monitored at a wavelength of 280 nm. An elution profile for the IgG4 solution at pH 7 was first obtained using a buffer solution containing 0.01 M sodium phosphate and 0.2 M NaCl. A strong peak at 13 cm^3^ of the elution volume (EV) was observed. To trace the time-dependent behavior of acid-stressed IgG4, three IgG4 mAb-B samples were prepared and analyzed. The first sample (t1) was prepared by diluting a 125 mg/mL IgG4 formulated solution 10-fold with 0.1 M glycine-HCl buffer solution (pH 2.0), resulting in a pH of 2.8. After filtration through a filter unit with a pore size of 0.22 μm (Merck Millipore Ltd., Cork, Ireland), a 50-μL aliquot of the IgG4 solution was loaded onto the column and eluted at a flow rate of 0.5 mL/min using a buffer solution containing 0.1 M glycine-HCl and 0.2 M NaCl (pH 2.0). The other two samples (t2 and t3) were prepared by diluting the 125 mg/mL IgG4 formulated solution 10-fold with 0.01 M sodium phosphate buffer solution (pH 7.4) and dialyzing it against 0.1 M glycine-HCl buffer solution (pH 2.0) using a membrane with a molecular weight cutoff of 12–14 kDa (Scienova GmbH, Jena, Germany) for 0.7 h at 4 °C. Following dialysis, one sample (t2) was analyzed without further incubation, while the other sample (t3) was incubated for an additional 2.3 h. After filtration, a 50-μL aliquot of each IgG4 solution was loaded onto the column and eluted at a flow rate of 0.2 mL/min using the same buffer solution (0.1 M glycine-HCl and 0.2 M NaCl, pH 2.0). The exposure duration to the acid solution for these samples, as indicated in Figure 2, was approximated at an elution volume of 14 cm^3^.

**Figure 2.**
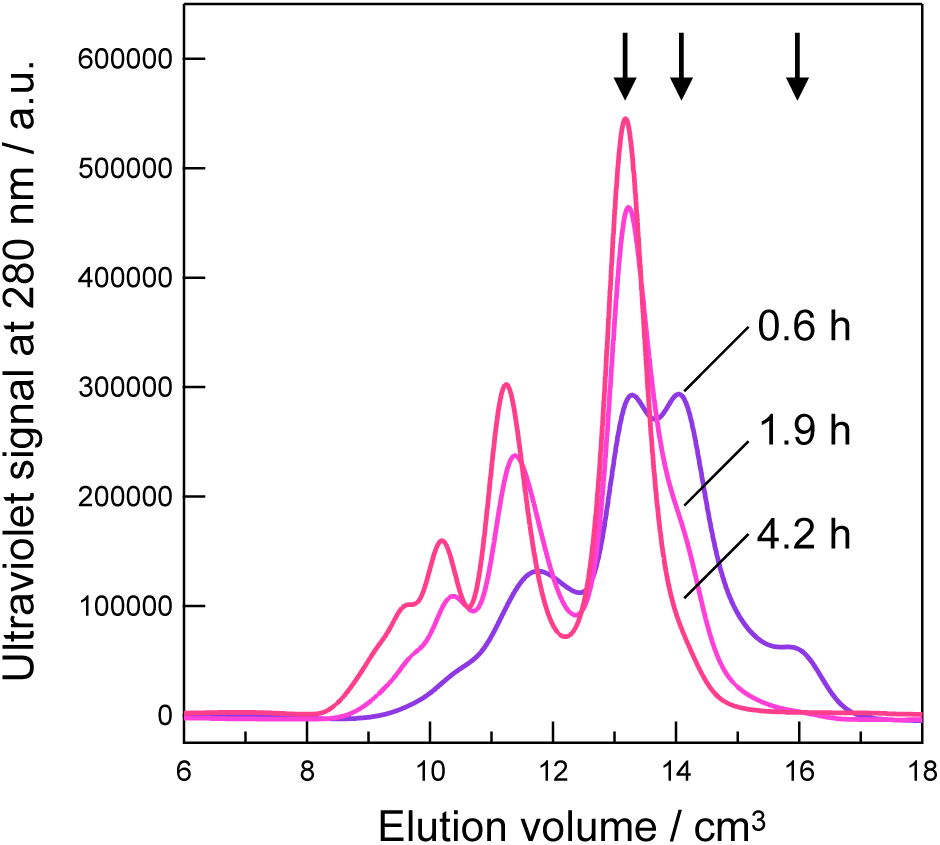
Size-Exclusion Chromatography Analysis of Acid-Stressed IgG4 mAb-B. Approximated exposure durations to the acid solution are indicated. The elution buffer solution consists of 0.1 M glycine-HCl and 0.2 M NaCl at pH 2.0.

For IgG1 mAb-A, the effect of brief exposure (approximately 30 minutes) to acidic conditions on SEC was also examined. The formulated IgG1 mAb-A powder was dissolved in 0.01 M sodium phosphate buffer solution (pH 7.4) and diluted 10-fold with 0.1 M glycine-HCl buffer solution (pH 2.0), resulting in a final concentration of approximately 10 mg/mL IgG1 mAb-A. After filtration, an 80-μL aliquot of the IgG1 solution was loaded onto the column and eluted at a flow rate of 0.5 mL/min using a buffer solution containing 0.1 M glycine-HCl and 0.2 M NaCl (pH 2.0).

### SAXS

Samples of IgG4 (1.2 mg/mL) at pH 7 (ID: 20241118ebg1) were prepared by dialysis against 0.01 M sodium phosphate buffer solution (pH 7.4). Acid-stressed IgG4 and IgG1 samples (ID: 20241118s1, 20241118s4, and 20230520s3) were prepared and analyzed using SEC-SAXS with a buffer solution containing 0.1 M glycine-HCl and 0.2 M NaCl (pH 2.0). The sample labeled 20241118s1 was prepared by diluting a 125 mg/mL IgG4 formulated solution 10-fold with 0.1 M glycine-HCl buffer solution (pH 2.0). After filtration, a 100-μL aliquot of the IgG4 solution was loaded onto the previously described Superdex200 column and eluted at a flow rate of 0.5 mL/min. This sample was acid-stressed for 0.7 h, corresponding to an elution volume of 14 cm^3^. The sample labeled 20241118s4 was prepared by diluting a 125 mg/mL IgG4 formulated solution 10-fold with 0.01 M sodium phosphate buffer solution (pH 7.4), followed by dialysis against 0.1 M glycine-HCl buffer solution (pH 2.0) for 1 h at 4 °C.

After dialysis, the sample was incubated for 0.6 h at 25 °C. After filtration, a 200-μL aliquot of the IgG4 solution was loaded onto the column and eluted at a flow rate of 0.2 mL/min. This sample was acid-stressed for 2.8 h at an elution volume of 14 cm^3^. For the sample labeled 20230520s3, a 10 mg/mL IgG1 solution in 0.01 M sodium phosphate buffer (pH 7) was dialyzed against 0.1 M glycine-HCl buffer solution (pH 2.0) for 3.9 h at temperatures ranging from 4 to 16 °C. After dialysis, the sample was incubated for 0.8 h. Following filtration, a 400-μL aliquot of the IgG1 solution was loaded onto the column and eluted at a flow rate of 0.2 mL/min. During the SEC-SAXS measurements, the protein concentrations were <1.2 mg/mL for IgG1 mAb-A and <1.0 mg/mL for IgG4 mAb-B, in which case the effects of interparticle interference on radius of gyration (*R*_g_) were within the uncertainty range and deemed negligible (Figure S1). These SAXS experiments provided valuable insights into the structural changes occurring in acid-stressed IgG4 and IgG1 samples, with the exposure times and conditions calibrated to achieve accurate assessments of molecular behavior.

The procedures for SAXS experiments and analyses were detailed in our previous paper[9]. Briefly, SAXS experiments were conducted on beamline BL-10C[14] at the Photon Factory of the High Energy Accelerator Research Organization (KEK) in Tsukuba, Japan. The X-ray wavelength (*λ*) was 0.15 nm, and the camera length was set to either 1 m or 2 m, calibrated using the scattering pattern of silver behenate. The scattering parameter *q* is defined as *q* = |***q***| = 4πsin*θ*/*λ*, where ***q*** is the scattering vector and 2*θ* is the scattering angle of the X-rays. X-ray intensities were recorded using a PILATUS3 2M detector (DECTRIS Ltd., Baden, Switzerland). For SEC-SAXS measurements, samples were passed through a stainless-steel sample cell via a Superdex 200^TM^ Increase 10/300 GL column operated by an HPLC system (Nexera-i; Shimadzu Co., Kyoto, Japan). The temperature of the sample cell and the column was maintained at 25.0 ± 0.1 °C. The exposure time was 20 s for SEC-SAXS measurements and 2 s for batch SAXS measurements. Circular 1D averaging of the scattering images was performed using the *Nika* program[15]. The absolute scattering intensity of the protein (*I*(*q*)) (cm^-1^) was determined using water scattering as a standard[16]. For dilute protein solutions, *I*(0)/*c* is rationalized by the molar mass (*M*) of the (a)glycosylated protein as:[9]

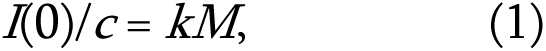

where *k* is defined as [(1 – *f*_g_)*v*(*ρ*_m_ – *ρ*_solv_) + *f*_g_*v*_g_(*ρ*_g_ – *ρ*_solv_)]^2^/*N*_A_. Here, *N*_A_ is Avogadro’s number, (*ρ*_m_ – *ρ*_solv_) is the electron density difference between the protein and the solvent (2.8 × 10^10^ cm^-2^)[17], and (*ρ*_g_ – *ρ*_solv_) is the electron density difference between the carbohydrate and the solvent (4.7 × 10^10^ cm^-2^)[18]. *f*_g_ is the weight fraction of the carbohydrate component (0.017 for mAb-A and 0.018 for mAb-B). *v* and *v*_g_ are the partial specific volumes of proteins (0.7425 cm^3^/g)[17] and carbohydrates (0.625 cm^3^/g)[18], respectively. Porod correction[9] was not applied to the present data.

CorMap analysis[19] for the SAXS data of acid-stressed IgG4 and IgG1 was conducted using the *DATCMP* tool in the ATSAS 3.0.4 software package[20]. The data were compared over the range *q* = 0.015–0.27 after appropriate scaling of the intensity. This scaling accounts for uncertainties, including those arising from the molar extinction coefficients of proteins. Unless otherwise stated, all analyses were performed using Igor Pro version 9.0.0 (WaveMetrics, Portland, OR). Details of the SAXS measurements and data analyses are provided in Table S1 (Supporting information).

### SAXS of IgG4 model structure

Since no published coordinates for IgG4 mAb-B were available, an IgG4 mAb-B model was constructed to calculate the SAXS profile. An initial model was generated using the homology modelling tool *MODELLER* 10.5[21], based on the IgG1 template bpj_1831_mmc2.pdb[22]. Disulfide bonds were specified according to information provided in the interview form[13]. The two heavy chains in the model were glycosylated at residue N295, showing a glycan structure Gal(β1-4)GlcNAc(β1-2)Man(α1-3)[Gal(β1-4)GlcNAc(β1-2)Man(α1-6)]Man(β1-4)GlcNAc(β1-4)[Fuc(α1-6)]GlcNAc(β1-(GlyTouCan ID: G78059CC) using the *Glycan Modeler* tool[23]. The IgG4 model was solvated using the *solution builder* in CHARMM-GUI[24], and the CHARMM36m force field was applied. The model structure was relaxed and equilibrated via molecular dynamics simulations using *GROMACS* 2021.4^14^ for 60 ns at 298.15 K and pH 7.0 in the presence of 0.15 M NaCl. Theoretical SAXS profiles were calculated based on the atomic coordinates of the IgG4 models using *CRYSOL*[25]. The model that best fit the experimental SAXS data (Figure 4) yielded an *R*_g_ value of 48.42 Å.

## Results and Discussion

Figure 2 presents the SEC data of acid-stressed IgG4 mAb-B for varying exposure durations (approximately 0.6, 1.9, and 4.2 hours; see Materials and Methods) at room temperature. Multiple peaks and shoulders are observed, corresponding to the monomer and aggregates. The contents of these aggregates eluted at < 12 cm^3^ elution volume (EV) as reported in our previous study[9]. The levels of these aggregates increased with prolonged exposure to pH 2. Notably, for brief acid exposure (0.6 h), three peaks appear at >13 cm^3^ EV, as indicated by the arrows in Figure 2. Based on the peak observed at 13 cm^3^ EV for native IgG4 mAb-B at pH 7 (data not shown) and previous literature[9], the peak at 13 cm^3^ EV is attributed to the monomer. Peaks at 14 cm^3^ and 16 cm^3^ EV diminished with extended acid exposure, suggesting they represent transient species. The subsequent SEC-SAXS experiments aim to investigate the origin of these peaks.

Figure 3 presents the SEC-SAXS results for 0.7 h and 2.8 h of exposure to an acidic solution (pH 2). The chromatograms for SAXS and ultraviolet (UV) absorbance at 280 nm are shown, along with *I*(0)/*c* and *R*_g_ values derived from each SAXS frame. *R*_g_, the radius of gyration, represents the size of the scattering particle, while *I*(0) is the absolute scattering intensity at *q* = 0, and *c* denotes protein concentration. The ratio *I*(0)/*c* is proportional to molar mass. At > 13 cm^3^ EV, *I*(0)/*c* remains constant at approximately 0.1 cm⁻¹ (mg/mL)⁻¹, corresponding to a molar mass of 1.4 × 10² kg/mol. This value aligns closely with 0.108 cm⁻¹ (mg/mL)⁻¹ for the theoretical molar mass of the monomer (1.48 × 10² kg/mol). Thus, all species eluting at > 13 cm^3^ EV are classified as monomers. The well-aligned SAXS-derived and UV-derived chromatograms in this region further support this assignment. In this study, the 148 kg/mol IgG4 molecule—comprising two heavy chains and two light chains —is referred to as a monomer. The peak at 11–12 cm^3^ EV corresponds to a dimer, as determined by its *I*(0)/*c* value. All three peaks at >13 cm^3^ EV represent IgG4 monomeric species separated by SEC, indicating conformational variation within the IgG4 monomer under brief exposure to acidic conditions (pH 2). These species are designated as M13, M14, and M16, where M (monomer) is followed by the elution volume. M16 and M14 are short-lived species at pH 2 (< approximately 1 h). To capture them, a flow rate of 0.5 cm^3^/min was necessary during the experiment to elute the 14–16 cm^3^ EV region within their lifetimes. Figure 2 demonstrates that for a 1.9 h exposure to the acidic solution (pH 2), M14 exhibits a longer lifetime than M16. Prolonged exposure leads to M13 becoming the dominant species. The molecular lifetimes at pH 2 can be ordered as M13 > M14 > M16. Notably, such intermediates were not detected for IgG1 mAb-A; brief exposure to acid did not yield observable peaks at the 14–16 cm^3^ EV region but produced a strong peak at 13 cm^3^ EV.

**Figure 3.**
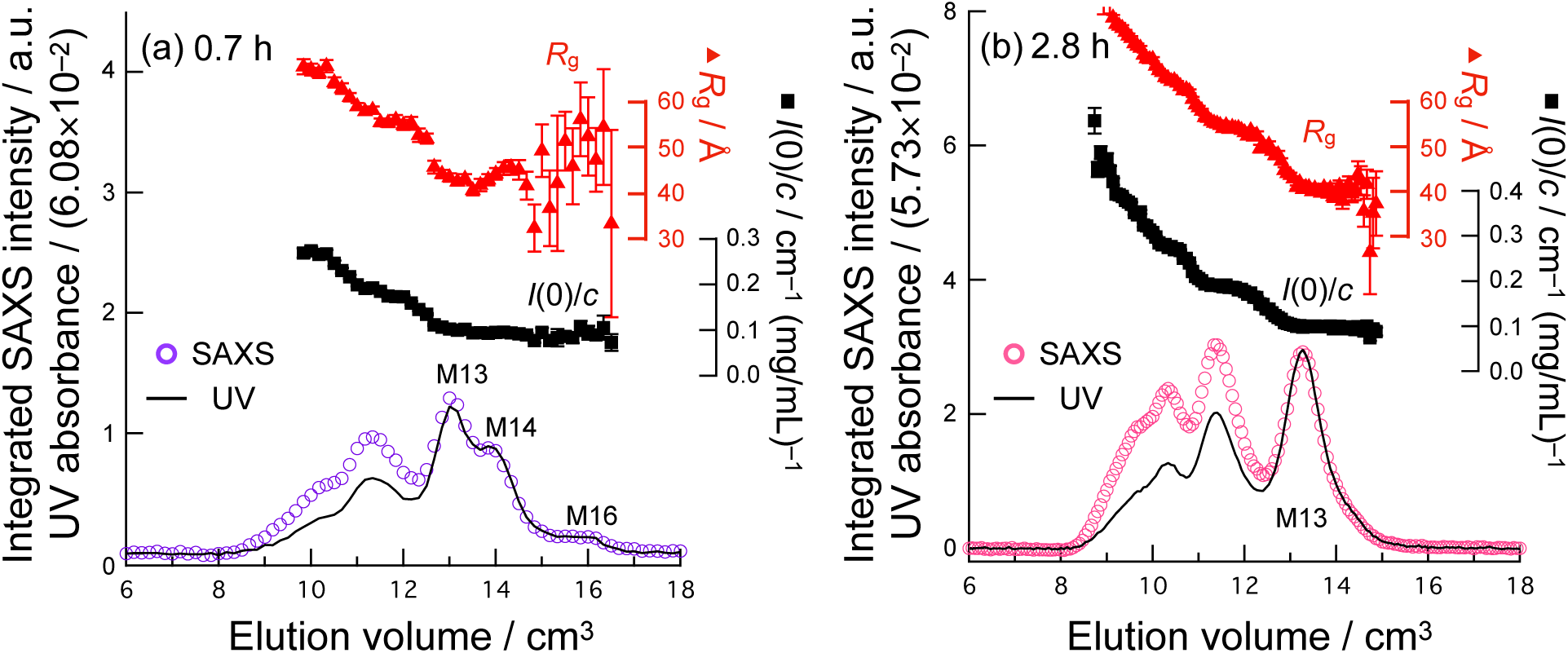
SEC-SAXS Analysis of IgG4 mAb-B. (a) 0.7 h and (b) 2.8 h exposure to an acid solution (pH 2) at 25 °C. The SAXS- and UV-derived elution profiles are overlaid, with the UV signals being scaled to match the SAXS signal at > 13 cm^3^ EV to effectively share the y-axis. The scaling factors are indicated in the labels of the *y*-axis. The SAXS-derived chromatogram represents integrated SAXS intensity versus retention time, where the integrated SAXS intensity is calculated by ΣI_raw_(*qi*), with I_raw_(*q*) representing the unprocessed raw scattering intensity and *q_i_* indicating the experimentally available *q*. This UV elution profiles were recorded simultaneously, capturing IgG4 mAb-B absorbance at 280 nm. The UV absorbance at 280 nm is independent of molar mass (molecular weight), while the integrated SAXS intensity depends on it. When both are scaled to the monomeric species, discrepancies between the profiles highlight differences in molar mass. Radius of gyration (*R_g_*) and *I*(0)/c values were determined for each frame.

The SAXS profiles of M13, M14, and M16 (Figure 4a) were extracted by averaging SAXS frames corresponding to 13.0–13.5, 14.0–14.5, and 15.5–16.5 cm^3^ EV, respectively (Figure 3a). *R*_g_ values were 42.8 ± 0.5 Å for M13, 45.1 ± 0.9 Å for M14, and 54.5 ± 6.3 Å for M16. The *R*_g_ values of M14 and M13 were smaller than the *R*_g_ of 48.3 ± 0.7 Å measured for native IgG4 mAb-B at pH 7. The SAXS profile of native IgG4 mAb-B was consistent with the theoretical profile derived from an IgG4 model (Figure 4a), showing characteristic peaks at *q*_1_, *q*_2_, and *q*_3_ in the Kratky plot (Figure 4b). These Kratky peaks were similar to those observed for native IgG1[28]. While M16 may represent residual native IgG4 species that are unstable at pH 2, the low signal-to-noise ratio limits definitive conclusions. M14 appears to be a transiently accumulated intermediate. The Kratky plots for M13 and M14 (Figure 4b) exhibit diminished *q*_2_ and *q*_3_ peaks compared to native IgG4, indicating a loss of native domain-domain correlations. The rapid elution of the smaller species observed in the present SEC experiment does not conform to the general rule of SEC, where larger species elute faster, assuming no protein-column interactions. This suggests that electrostatic and/or hydrophobic interactions between the column and protein molecules[29] play a significant role. The degree of structural organization of the IgG4 monomer is presumed to be coupled with protein-column interactions and elution volume (EV). The molecular lifetimes, *R*_g_ values, and elution volumes suggest a sequential pathway for acid-denaturation of IgG4: M16 ® M14 ® M13.

**Figure 4.**
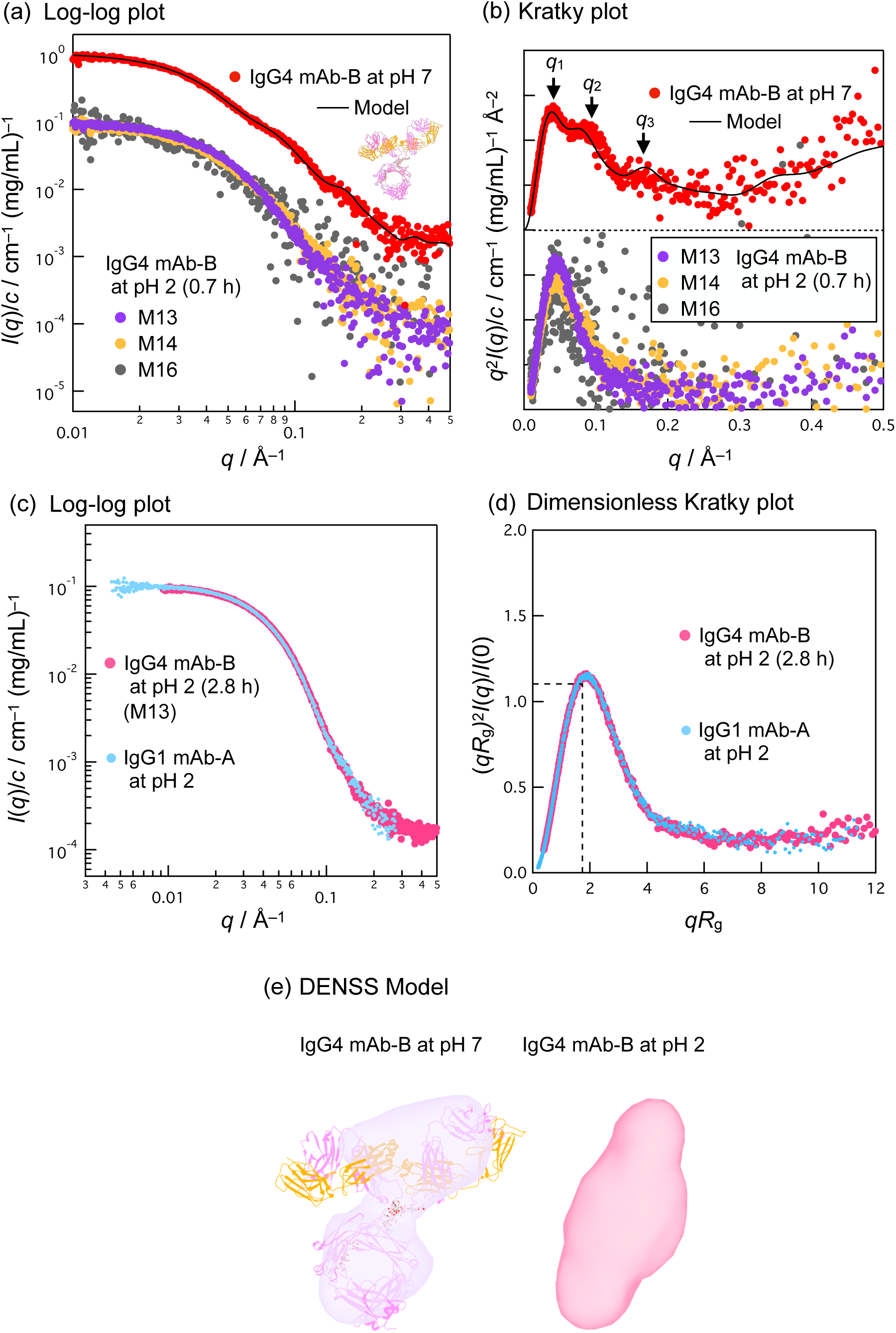
SAXS Profiles of IgG4 mAb-B. (a) Log-log plot and (b) Kratky plot at pH 2 (0.7 h exposure) and pH 7. The “Model” line represents the theoretical SAXS profile calculated from the atomic coordinates of an IgG4 mAb-B model (see Materials and Methods). (c-e) SAXS of IgG4 mAb-B at pH 2 (2.8 h exposure) and comparison with IgG1 mAb-A SAXS at pH 2: (c) Log-log plot and (d) dimensionless Kratky plot. Well-folded proteins display a local maximum at (1.732, 1.104) (black dashed lines)[26]. (e) *Ab initio* models generated by the DENSS method[27].

Because the monomer species M13 is long-lived and stable at pH 2, SAXS data with an improved signal-to-noise ratio (Figure 4c and 4d) were obtained by using longer X-ray exposure times and a slower flow rate (0.2 cm^3^/min). The *R*_g_ of M13, determined at 13.0–13.5 cm^3^ EV in Figure 3b, was 41.3 ± 0.1 Å, comparable to the *R*_g_ values of 40.5 ± 0.4 Å[9] and 41.8 ± 0.2 Å (the present study) reported for the acid-denatured structure of IgG1. Both IgG1 and IgG4 become smaller upon acid denaturation than their native forms; this is one of the primary findings of the present study. Superimposing the SAXS profiles of IgG1 and IgG4 reveals that they are almost identical (Figure 4c and 4d). The hypothesis of similarity between IgG4 M13 and IgG1 AFS SAXS data could not be rejected (CorMap test;[19], 351 points, *C* = 15, *P* = 0.01), where a significance level of 0.01 or less is preferred[19]. While the SAXS profiles of their native structures (pH 7) differ significantly (as shown in this study and reported in previous studies[11,12]), their acid-denatured AFS structures are identical. The shapes of native and AFS IgG4 were reconstructed by the DENSS method[27] (Figure 4e). It is to be noted that while the DENSS model does not exclude the possibility of multiple conformations fitting the SAXS profile, it highlights the dominant structures.

The common AFS formation observed in both IgG1 and IgG4 strengthens the hypothesis that AFS is not restricted to specific antibodies. This study compared IgG1 and IgG4, which have different variable region sequences, and primarily supports previous findings that variations in the variable region do not influence AFS formation[9]. In addition, the minor differences in the constant region sequences between IgG1 and IgG4 appear to have little impact. Despite differences in hinge region length and disulfide bond arrangement (Figure 1), in the present study, we demonstrate that the anomalous compaction associated with AFS formation is resilient to these structural variations. As described in our previous paper [9], acid-induced compaction requires “intramolecular aggregation” involving domain reorientation, where non-native Fc-Fab interactions stabilize the AFS. Further, in the present study, we revealed that the common residues between IgG1 and IgG4, particularly in constant regions, play a key role in stabilizing the AFS. Variable regions, such as complementarity-determining regions, may not be involved in non-native contacts within the AFS.

Compactability and size reduction through acid denaturation are common among the studied antibodies. While the identical SAXS profiles displayed for the IgG1 and IgG4 AFSs are noteworthy, we have reservations regarding whether AFS antibodies can yield a single SAXS profile or *R*_g_, considering that the AFS is plastic [9]. For example, the *R*_g_ of IgG1 in the previous study [9] is dependent on the salt (NaCl) concentration and/or the mAb species. Salt provides a shielding effect against intramolecular electrostatic repulsion, which can vary depending on nature of the antibody used.

Circular dichroism (CD) spectra were measured to monitor secondary structural changes during acid denaturation (Figure S2). Common spectral changes induced by acid were identified among IgG4, previously studied IgG1s[3,4,9], and antibody fragments (Fab[30] and C_H_3[31]): Specifically, ellipticities in the ∼200 nm region were positive (convex upward curves) at pH 7 but inverted to negative (convex downward curves) at pH 2. Spectral decomposition of the CD data using BestSel[32] indicates that IgG4 mAb-B at pH 2 retains a *β*-rich structure (42.5%: antiparallel *β* [31.6%] and parallel *β* [10.9%]), comparable to its *β*-rich content at pH 7 (48.6%: antiparallel *β* [44.0%] and parallel *β* [4.6%]). This rules out the complete unfolding of the secondary structure, instead suggesting a minor reorganization of the local or secondary structures. These findings align with the case of IgG1 AFS, where minor alterations to *β*-rich structures have been confirmed by CD and infrared spectroscopy[9].

In summary, despite differences in sequence and higher-order structure between humanized IgG1 and IgG4, both monomeric species are capable of forming the AFS structure under acidic conditions, with the AFS structures being identically compact. This study rejects the notion that AFS formation in IgGs is sequence-or subclass-specific. The acid-induced compaction associated with AFS appears to be a widespread phenomenon among IgGs. Furthermore, during acid denaturation, intermediate structure(s) were observed for IgG4 mAb-B, which could be separated by SEC. These intermediates were not detected for IgG1 mAb-A. The IgG4 format, including its disulfide bonds, may confer a longer molecular lifetime of the intermediate(s) during AFS formation. These findings provide valuable insights into the common mechanisms underlying denaturation and aggregation of therapeutic IgG1 and IgG4 antibodies and can inform further advancements in the development of antibody drugs.

## Author Contributions

Conceptualization, H. I. and S. H.; methodology, H. I.; software, H. I.; validation, H. I. and S. H.; formal analysis, H. I.; investigation, H. I. and S. H.; data curation, H. I.; writing—original draft preparation, H. I.; writing—review and editing, S. H.; visualization, H. I.; funding acquisition, H. I. and S. H. All authors have read and agreed to the published version of the manuscript.

## Abbreviations

IgG: immunoglobulin G
SAXS: small-angle X-ray scattering
SEC: size-exclusion chromatography
EV: elution volume
CD: circular dichroism.

## Acknowledgements

This research is partially supported by the Japan Society for the Promotion of Science [Grants numbers JP21K06503 (to H.I.) and JP22K06575 (to S.H.)]. The SAXS experiment was performed under the approval of the Photon Factory Program Advisory Committee (Proposals 2024G067). Computations were partially performed on the NIG supercomputer at the ROIS National Institute of Genetics. The authors declare no competing financial interest. The authors are grateful to Dr. Risa Shibuya (Osaka University, Suita, Japan) for the Graphical Abstract illustration.

## Data availability

The data that support the findings of this study are openly available in the Small Angle Scattering Biological Data Bank (SASBDB) as entries SASDW64 (https://www.sasbdb.org/data/ SASDW64), SASDW54 (https://www.sasbdb.org/data/SASDW54), and SASDW44 (https://www.sasbdb.org/data/SASDW44). Additional materials, methods, and results of SEC-SAXS and CD are available in the supporting informationl of this article.

