## Supplementary Information for "Alternatively Folded State of IgG4: Acid-Induced Compaction is Common between IgG1 and IgG4"

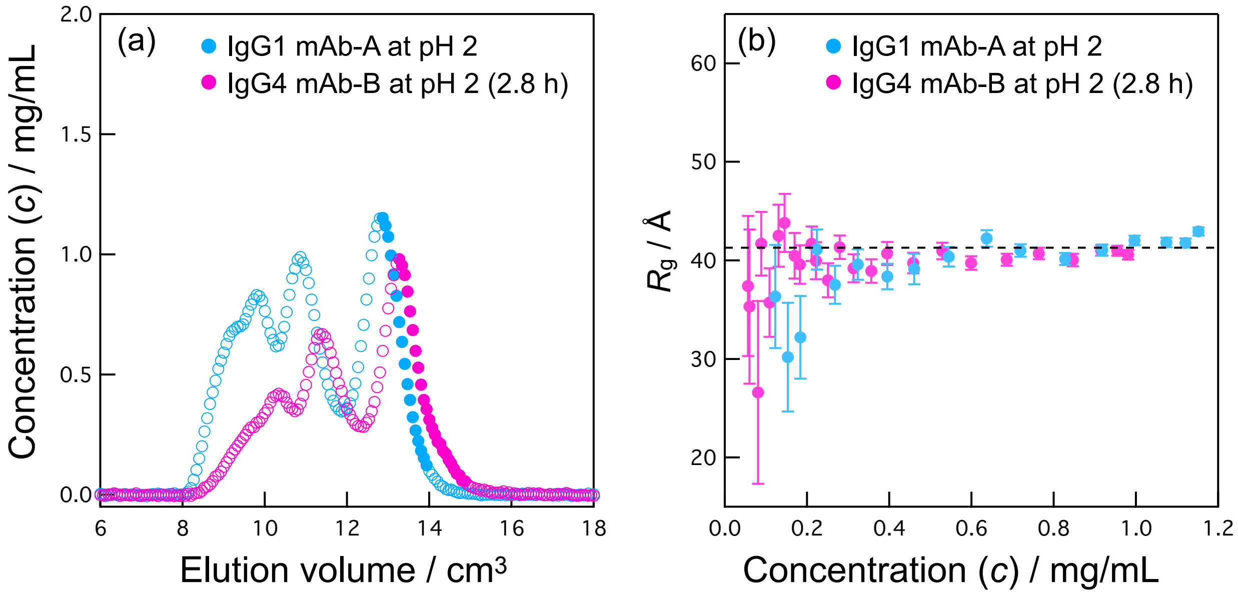


Figure S1. Protein concentration data during SAXS-monitored SEC elution to evaluate interparticle interference effects. (a) Protein concentration data spanning the SEC time course, with the *x*-axis showing the elution volume (0.2 mL/min flow rate). (b) Concentration-dependence of $R_g$ of IgG1 mAb-A and IgG4 mAb-B at pH 2. The data points presented in (a) as filled circles were used. The dashed line indicates 41.3 Å.

**CD measurements** A 125 mg/mL IgG4 mAb-B formulated solution was diluted 10-fold with 0.01 M sodium phosphate buffer solution (pH 7.4) and dialyzed against the same phosphate buffer solution for 3 h at 4 °C to prepare the mother solution. The native IgG4 sample was then prepared by diluting the mother solution 100-fold with the phosphate buffer solution. For the acid-denatured IgG4 sample, the mother solution was similarly diluted 100-fold but with 0.1 M glycine-HCl buffer solution (pH 2.0), followed by dialysis against a buffer solution containing 0.1 M glycine-HCl and 0.2 M NaCl (pH 2.0) for 0.7 h at 25 °C CD spectra were recorded using a J-820 CD spectrometer (JASCO Co. Ltd., Tokyo, Japan) equipped with a temperature-controlled cell with a 0.1-cm path length, maintained at 25 °C. The raw ellipticities were converted to mean residue ellipticities using the equation $[\theta] = \theta_{obs}/(10nc_Ml)$, where $\theta_{obs}$ is the observed ellipticity (millidegrees), $l$ is the optical path length (cm), $c_M$ is the molar concentration of the protein, and $n$ is the number of residues.

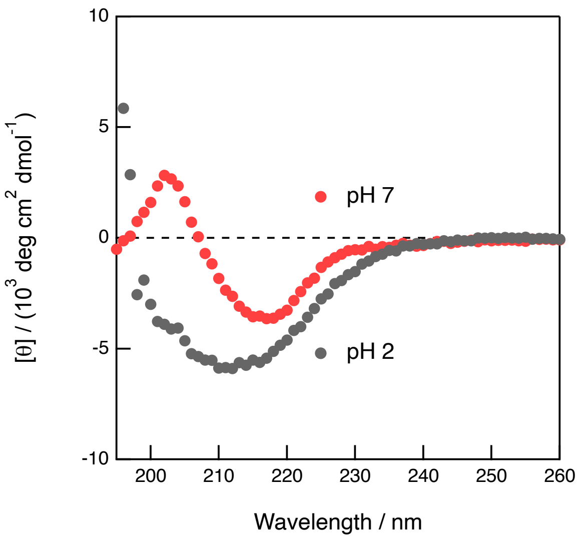


Figure S2. Circular dichroism (CD) analysis. The CD spectra of native IgG4 mAb-B (pH 7, red) and acid-denatured IgG4 mAb-B (pH 2, grey) are shown.

**Table S1.** SAXS sample details, data collection, analysis, and 3D modelling details.

| (*a*) Sample details | | | | |
|---|---|---|---|---|
| Organism | *Mouse/human* | | | |
| Source | Chinese hamster ovary cell recombinant expression | | | |
| Sample ID | 20241118ebg1 | 20241118s1 | 20241118s4 | 20230520s3 |
| *Scattering particle composition* | IgG4 mAb-B | | | IgG1 mAb-A |
| Protein(s) | IgG4 mAb-B [DrugBank DB11914] | | | IgG1 mAb-A [DrugBank DB00072] |
| Carbohydrates/glycans | [GlyTouCan ID: G80858MF + $Gal_{0-2}$][a] | | | [GlyTouCan ID: G39213VZ, G80858MF, G89993FE, G27919IH, G58667NI, G78059CC][b] |
| *Sample environment/configuration* | | | | |
| Solvent composition | 0.01 M sodium phosphate, pH 7.4 | 0.1 M glycine-HCl, 0.2 M NaCl, pH 2.0 | | |
| Sample temperature (°C) | 25.0 ± 0.1 °C | | | |
| In beam sample cell | 1 mm path length with quartz window, no flow | 1 mm path length with quartz window, flow | | |
| *Batch measurements* | | | | |
| Sample concentration(s), mg/ml | 1.2 | | | |
| *Size Exclusion Chromatography SEC-SAXS* | | | | |
| Sample injection concentration, mg/ml | | ~12.5 | ~12.5 | ~10 |
| Sample injection volume, mL | | 0.1 | 0.2 | 0.4 |
| SEC column type | | Superdex200™ Increase (10/300) GL (Lot. 10299331) | | |
| SEC flowrate, mL/min | | 0.5 | 0.2 | 0.2 |

[a] According to the interview form[8].

[b] Based on information in PMDA (Pharmaceuticals and Medical Devices Agency): Review report regarding the anti-HER2 humanized monoclonal antibody Herceptin® Injection 60 and 150, dated January 16, 2008 (https://www.pmda.go.jp/drugs/2007/P200700066/45004500_21300AMY00128_A100_2.pdf).

**Table S1 (continued).**

| (*b*) SAS data collection | | | | |
|---|---|---|---|---|
| Data acquisition/reduction software | PILATUS Measurement Control Software at Photon Factory/*Nika*[1] | | | |
| Source/instrument description | Beamline BL-10C with Dectris PILATUS 2M detector, the Photon Factory (PF) of the High Energy Acceleration Research Organization (KEK)[2] | | | |
| Measured $q$-range ($q_{min} - q_{max}$; Å$^{-1}$) | 0.01–0.56 | | | 0.004–0.27 |
| Method for scaling intensities | Absolute scaling (cm$^{-1}$) referenced to water | | | |
| Exposure time(s), number of exposures. For SEC-SAS, final number of sample frames used for averaging. | 2 s × 5 frames | M13&M14: 20 s × 4 frames<br>M16: 20 s × 7 frames | 20 s × 9 frames | 20 s × 13 frames |
| Additional relevant details | 1078 mm sample-to-detector distance | | | 2080 mm sample-to-detector distance |

**Table S1 (continued).**

| (*c*) SAS-derived structural parameters | | | | |
|---|---|---|---|---|
| Methods/Software | *Igor Pro* (9.0.0) and *GNOM* (ATSAS 3.0.4)[3] . | | | |
| *Guinier Analysis* | 20241118ebg1 | 20241118s1 | 20241118s4 | 20230520s3 |
| $I(0)/c \pm \sigma$ ($cm^{-1}$ $(mg/mL)^{-1}$) | 0.11 ± 0.001 | M13: 0.10 ± 0.001<br>M14: 0.10 ± 0.001<br>M16: 0.11 ± 0.01 | M13: 0.10 ± 0.0002 | 0.10 ± 0.0001 |
| $R_g$ ± σ (Å) | 48.3 ± 0.7 | M13: 42.8 ± 0.5<br>M14: 45.1 ± 0.9<br>M16: 54.5 ± 6.3 | M13: 41.3 ± 0.1 | 41.8 ± 0.2 |
| $min < qR_g < max$ limit | 0.72–1.30 | M13: 0.64–1.29<br>M14: 0.68–1.30<br>M16: 0.82–1.30 | M13: 0.62–1.30 | 0.31–1.29 |
| Linear fit assessment ($R^2$) | 0.94 | M13:0.94<br>M14:0.88<br>M16: 0.25 | M13: 1.00 | 0.99 |
| *PDDF/P(r) analysis* | 20241118ebg1 | 20241118s1 | 20241118s4 | 20230520s3 |
| $I(0) \pm \sigma$ ($cm^{-1}$ $(mg/mL)^{-1}$) | 0.11 ± 0.001 | M13: 0.10 ± 0.0004<br>M14: 0.09 ± 0.0006<br>M16: 0.09 ± 0.0005 | M13: 0.10 ± 0.0002 | 0.09 ± 0.0001 |
| $R_g$ ± σ (Å) | 48.8 ± 0.3 | M13: 41.8 ± 0.2<br>M14: 43.5 ± 0.3<br>M16: 48.9 ± 0.2 | M13: 41.6 ± 0.1 | 41.7 ± 0.1 |
| $d_{max}$ (Å) | 156 | M13: 127<br>M14: 132<br>M16: 144 | M13: 140 | 129 |
| $q$-range ($Å^{-1}$) | 0.015–0.166 | M13: 0.015–0.186<br>M14: 0.015–0.177<br>M16: 0.015–0.073 | M13: 0.015–0.194 | 0.007–0.191 |
| $P(r)$ fit assessment<br>(Total estimate from *GNOM*) | 0.90 | M13: 0.80<br>M14: 0.88<br>M16: 0.60 | M13: 0.82 | 0.83 |

**Table S1 (continued).**

| (*d*) Scattering particle size | | | | |
|---|---|---|---|---|
| Methods/Software | *PRIMUS/qt* (ATSAS 3.0)[3] for *M* from Bayesian inference[4] and $V_P$, equation (1) in this paper for *M* from $I(0)/c$ | | | |
| | 20241118ebg1 | 20241118s1 | 20241118s4 | 20230520s3 |
| *Volume estimates* | | | | |
| Porod volume, $V_P$ ($10^3$ Å$^3$) (ratio to M) | 211 (1.4) | M13: 246 (1.7)<br>M14: 218 (1.5)<br>M16: 358 (2.4) | 238 (1.6) | 240 (1.6) |
| *Molecular weight (M) estimates (kDa)* | | | | |
| From chemical composition | | 148 | | 148 |
| From SAS, concentration independent method (Bayesian inference, range with % confidence) | 127–151, 90% | M13: 163–195, 98%<br>M14: 151–195, 97%<br>M16: 195–264, 99% | M13: 134–177, 97% | 163–195, 99% |
| From $I(0)/c$ (ratio to expected) | 146 ± 2 (0.99) | M13: 139 ± 1 (0.94)<br>M14: 132 ± 2 (0.89)<br>M16: 147 ± 13 (0.99) | M13: 141 ± 0.3 (0.95) | 131 ± 0.2 (0.89) |
| Partial specific volume, (cm$^3$/g) | 0.7425 for the polypeptides[5]<br>0.625 for the carbohydrates[6] | | | |
| Contrast, $\Delta\rho$ ($10^{10}$ cm$^{-2}$) | 2.8 for the polypeptides[5]<br>4.7 for the carbohydrates[6] | | | |
| From SAS-independent measure | n.a. | n.a. | n.a. | n.a. |

**Table S1 (continued).**

| (*e*) Modelling | | | |
|---|---|---|---|
| *Shape modelling method(s)/Software* | *Ab initio* (*DENSS*)[7] | *Ab initio* (*DENSS*)[7] | |
| | 20241118ebg1 | 20241118s4 | |
| Software | *DENSS* 1.7.4 | *DENSS* 1.7.4 | |
| $q$-range for fit ($q_{min} - q_{max}$; Å$^{-1}$) | 0.015–0.166 | 0.015–0.194 | |
| Symmetry/anisotropy assumptions | No | No | |
| Number of individual model reconstructions | 100 | 100 | |
| CorMap *P*-values for fit | Avg. 0.57 | Avg. 0.26 | |
| (*f*) Data and model deposition | | | |
| | 20241118ebg1 | 20241118s4 | 20230520s3 |
| SASBDB IDs | SASDW64 | SASDW44 | SASDW54 |
